# Contribution of *rsaC*, a small non-coding RNA, towards the pathogenicity of *Staphylococcus aureus* in a mouse systemic infection model

**DOI:** 10.1101/2021.10.19.465069

**Authors:** Suresh Panthee, Hiroshi Hamamoto, Atmika Paudel, Suguru Ohgi, Kazuhisa Sekimizu

**Affiliations:** Drug Discoveries by Silkworm Models, Faculty of Pharma-Science, Teikyo University, Tokyo, Japan; Teikyo University Institute of Medical Mycology, Tokyo, Japan; Division of Sport and Health Science, Graduate School of Medical Care and Technology, Teikyo University, Tokyo, Japan; International Institute for Zoonosis Control, Hokkaido University, Sapporo, Japan; Laboratory of Microbiology, Graduate School of Pharmaceutical Sciences, The University of Tokyo, 7-3-1 Hongo, Bunkyo-ku, Tokyo 111-0033, Japan; Genome Pharmaceuticals Institute, Ltd, Tokyo, Japan; Kyowa Kirin Co., Ltd., 1-9-2 Otemachi, Chiyoda-ku, Tokyo 100-0004, Japan

**Keywords:** *Staphylococcus aureus*, Transcriptome analysis, virulence, sRNA, *in vivo*, pathogenicity

## Abstract

Understanding how a pathogen responds to the host stimuli and succeeds in causing disease is crucial for developing a novel treatment approach against the pathogen. Transcriptomic analysis facilitated by RNA-Seq technologies is used to examine bacterial responses at the global level. However, the ability to understand pathogen behavior inside the host tissues is hindered by much lower pathogen biomass than host tissue. Recently, we succeeded in establishing a method to enrich *Staphylococcus aureus* cells from infected organs. In this research, we analyzed the small non-coding RNA (sRNA) transcriptome of *S. aureus* inside the host and found that *rsaC* was among the highly expressed sRNAs. Furthermore, by gene disruption and complementation, we demonstrated that *rsaC* was required for full pathogenicity of *S. aureus* in a murine model. Besides, we found that Δ*rsaC* showed a difference in gene expression depending on the oxygen and host stress. The findings of this study suggest *rsaC* acts as a novel virulence factor in *S. aureus* and might facilitate the adaptation of *staphylococci* within the host.

**Importance:** Drug-resistant *Staphylococcus aureus* is among the pathogen for which new treatment options are urgently needed. However, limited understanding of *S. aureus* pathogenesis in the host has hindered unearthing potential strategies to treat the infections. Here, based on the *in vivo* transcriptomic analysis, we present the identification of a small non-coding RNA (sRNA) *rsaC* as a novel virulence factor of *S. aureus*. Furthermore, we performed transcriptomic analysis of the *rsaC* disrupted mutant and identified different pathways, possibly controlled by *rsaC*, during aerobic, anaerobic, and *in vivo* conditions. These findings contribute to reveal the role of sRNA *rsaC* and broadens our understanding of the adaptation of *S. aureus* to host environments.

## Text

The growth and behavior of a pathogenic microorganism differ between the host environment and in the *in vitro* culture. Understanding how a pathogen responds to the host stimuli and succeeds in causing disease is crucial for developing novel drugs against the pathogen. Comprehensive transcriptomic analysis facilitated by RNA-Seq technologies using next-generation sequencers has been widely used to understand pathogen response at the global level (1, 2). Several studies have used *in vitro* host-like or *in vivo* host environments to understand the infection process and alterations in the pathogen and the host (3, 4). Such studies aimed to elucidate the functional role of protein-coding genes, while the small non-coding RNAs (sRNAs) remain largely unattended. Studies that performed the comparative analysis of pathogen response in the host to the in vitro growth allowed us to evaluate how a pathogen behaves during infection situations and how the host responds to pathogen invasion (4–7). The current understanding of bacterial pathogenesis allowed us to identify key virulence determinants that could potentially be exploited as a drug target for antivirulence drugs. Whereas it can be expected that pathogens lacking these virulence factors are easy to be killed or cleared from the host, their behavior in the host has not been analyzed at the global level. Furthermore, a detailed evaluation of pathogenesis requires an understanding of pathogen behavior during actual infection conditions.

sRNAs have been identified in living organisms, including the bacterial kingdom. Although they do not encode functional proteins, they can accomplish a wide range of biological functions, including regulation of gene expression at the levels of transcription, RNA processing, mRNA stability, and translation (8). In bacteria, sRNAs have the potential to regulate the gene expression pattern of the bacteria by interacting with protein or mRNAs by cis- or trans-acting mechanisms (8), thus affecting multiple cellular processes such as pathogenesis and bacterial physiology in response to environmental cues, facilitating survival under unfavorable environments. In *Staphylococcus aureus*, a pathogen of global concern, several sRNAs have been identified (9–11), while the functional characterization of this class of RNAs has long been forsaken. One of the functionally characterized and extensively studied Staphylococcal sRNA, RNAIII, regulates the expression of several virulence factors both positively and negatively (12–14). *rsaC*, a part of polycistronic operon *mntABC*, is another staphylococcal sRNA known to be expressed in infected host (15) and modulate oxidative stress during manganese starvation (16). Whereas these studies provided a hint towards its role in *S. aureus* virulence, a cell-based study showed that the Δ*rsaC* strain was more persistent in macrophages and resistant to opsonophagocytosis than the wild-type strain (16) indicating its obscure role in pathogenesis.

We recently successfully enriched pathogen RNA from an infected animal and performed an *in vivo* RNA-Seq analysis of *Streptococcus pyogenes* (5) and *S. aureus* (6) using a two-step cell crush method. In this manuscript, we performed the comprehensive analysis of *S. aureus* sRNAs expressed within the host. Using RNA-Seq analysis of the *rsaC* disrupted mutant, we showed a clear difference between the gene expression patterns during *in vivo* and *in vitro* (aerobic and anaerobic) growth conditions. Furthermore, we found that the transcription of fermentation and oxidoreductase-related genes, virulence and toxin-related genes, and host evasion-related genes were affected during aerobic, anaerobic, and *in vivo* growth, respectively. Our results highlight the importance of *in vivo* transcriptomic analyses of two strains with the same genetic background to allow a direct comparison, revealing changes in different environments.

## Materials and Methods

### Ethics statement

All mouse experiments were performed at the University of Tokyo and Teikyo University, following the animal care and use regulations approved by the Animal Use Committee (approval numbers: P27-4 and 16-014 at respective institutes).

### Bacterial strains and primers used in the study

Bacterial strains and primers used in this study are listed in **Table 1**. *S. aureus* and *Escherichia coli* were routinely grown in LB or TSB medium, respectively, at 37°C with shaking. The media were supplemented with antibiotics, as required.

**Table 1:**
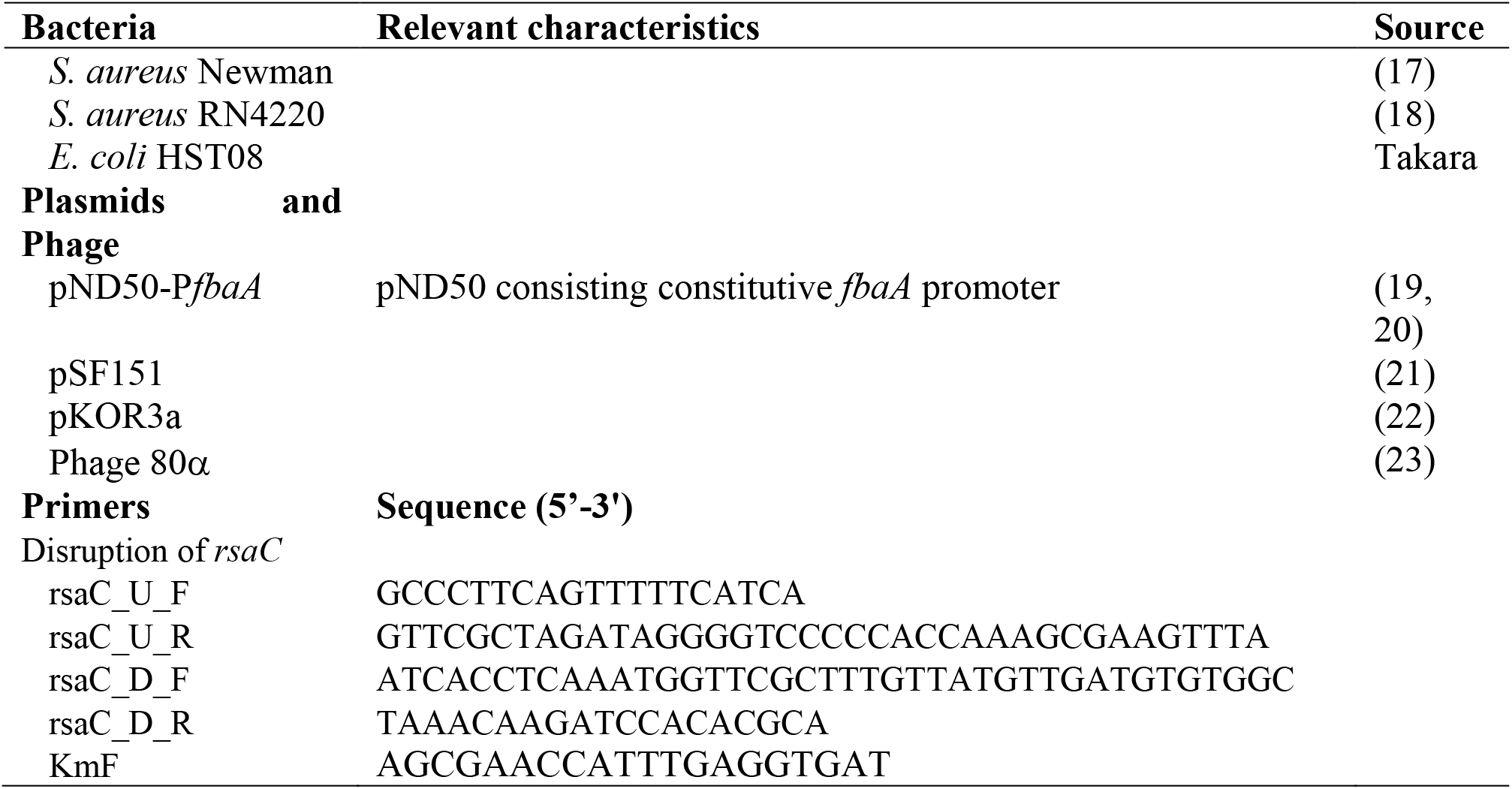
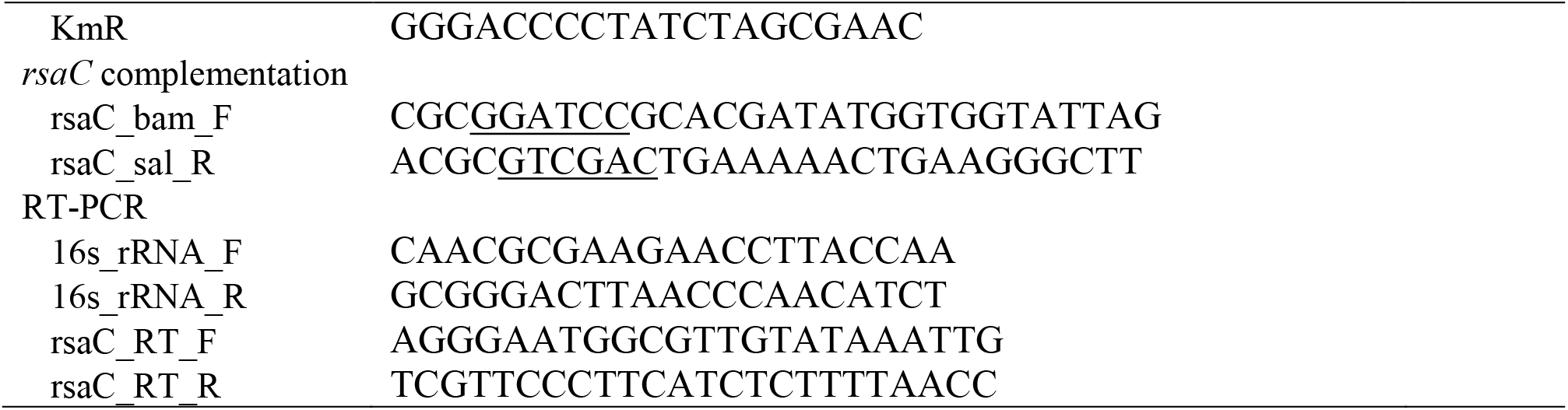
Bacterial strains, plasmids, and primers used in the study.

### Real-time RT-PCR

According to the manufacturer’s recommendations, one microgram of RNA was used to prepare cDNA using a High-Capacity RNA-to-cDNA™ Kit (Applied Biosystems; Foster City, CA, USA). From this, 15 ng of the cDNA was used as a template for RT-PCR using Fast SYBR Green Master Mix (Thermo Fisher Scientific) on a 7500 Fast (Applied Biosystems) machine with 40 cycles of denaturing at 95 °C followed by annealing/extension at 60 °C.

### Construction of Δ*rsaC* strain and complementation

*rsaC* was disrupted using a double cross-over recombination method as previously described (24). Briefly, the genome regions up-and down-stream of *rsaC* were amplified by PCR using the listed primers. Then, overlap extension-PCR was performed using these two DNA fragments together with the *aph* gene amplified from the pSF151 vector using primers KmF and KmR. The PCR product was cloned into the pKOR3a vector (24) and introduced into the RN4220 strain (18) by electroporation. Integration of the mutant cassette in the genome was confirmed by PCR and further transformed into *S. aureus* Newman (17) by phage transduction using phage 80α as previously described (23, 25).

To prepare the overexpression strain, the *rsaC* coding region was amplified by the primer pair rsaC_bam_F and rsaC_sal_R (**Table 1**) and ligated to BamHI-SalI digested pND50-P*fbaA* (19, 20). The resulting plasmid, pND50-P*fbaA*-*rsaC*, was then transformed into *S. aureus* RN4220 by electroporation and then to *S. aureus* Newman wild and Δ*rsaC* strains by phage transduction.

### Proteolytic and hemolytic assays

Bacteria were grown overnight in TSB medium, supplemented with antibiotics, as required, at 37°C with shaking. From this, 2 μl aliquots were spotted on TSB containing 3.3% skim milk or sheep blood agar plates (E-MR93; Eiken Chemical, Tokyo, Japan) to determine proteolysis and hemolysis, respectively. The plates were incubated at 37°C overnight. Sealed container with Anaero Pack (Mitsubishi Gas Chemical, Tokyo, Japan) were used to determine phenotypes under anaerobic conditions. The proteolytic and hemolytic activities were determined by the appearance of a clear zone surrounding the bacterial growth.

### Mouse survival assay

*S. aureus* Newman wild-type and Δ*rsaC* strains were grown overnight on TSB medium supplemented with antibiotics, as required, on a rotary shaker maintained at 37 °C to obtain full growth. The full growth was diluted 100-fold with TSB and cultured overnight on the same shaker, and then the cells were centrifuged and resuspended in phosphate-buffered saline (PBS, pH 7.2) to have an optical density (OD_600_) of 0.7. An aliquot (200 μl) of the prepared cell suspension was then injected intravenously into C57BL/6J mouse, and mouse survival was observed. Survival analysis was performed using GraphPad Prism ver 8.0 (GraphPad Software), and statistical analysis was performed using the Log-rank (Mantel-Cox) test.

### *S. aureus* infection, organ isolation and RNA isolation

*Staphylococci* were grown overnight on TSB medium at 37°C with shaking. The full growth was diluted 100-fold with 5 ml TSB and regrown for 16h under the same conditions. The cells were centrifuged and suspended in PBS (pH 7.2). The cells (OD_600_ = 0.7) were injected into C57BL/6J mice via the tail vein. On day 1, mice were killed to harvest organs. Organs were immediately placed either in ice to calculate viable cell numbers in each organ or liquid nitrogen and maintained at −80 °C for RNA extraction. Each experiment was conducted with three animals, and data are represented as an average. Mouse organs were homogenized, and total RNA was extracted, as explained previously (6). RNA extraction from *in vitro* culture was performed as explained (19). For anaerobic RNA-Seq, staphylococci were first grown aerobically to reach OD_600_ of 1.0 and then transferred to anaerobic growth for 30 minutes.

### Library preparation and RNA-sequencing

Total RNA was subjected to rRNA depletion using a MICROBExpress™ Kit (Thermo Fisher Scientific, Waltham, MA) and used for library preparation for RNA-Seq using an Ion Total RNA-Seq Kit v2 following the manufacturer’s instructions. After confirming the size distribution and yield of the amplified library using a bioanalyzer, the libraries were enriched in an Ion PI Chip v2 using the Ion Chef (Thermo Fisher Scientific) and sequenced using Ion Proton System (Thermo Fisher Scientific). These data have been deposited in the NCBI Sequence Read Archive under accession number ######.

### Differential gene expression analysis

All data were analyzed using CLC Genomics Workbench software, version 21.0.4 (CLC Bio, Aarhus, Denmark). Reads were aligned to the Newman genome annotated with ncRNA genes allowing a minimum length fraction of 0.95 and a minimum similarity fraction of 0.95. Differential gene expression analysis was performed using the default setting. Genes with a false discovery rate (FDR) *p*<0.05 were classified as having significantly different expressions.

## Results

### Comprehensive analysis of sRNAs expression in the host environment

To quantitatively analyze the expression of *S. aureus* sRNAs within the host, we first annotated the sRNAs present in the Newman strain based on the report of Sassai *et al*. (11). Next, we performed an RNA-Seq analysis based on the reads obtained from our two-step cell crush method (6). A comparative expression analysis of RNA isolated from host liver at 6- hr, 1- day and 2- day post-infection to the RNA isolated from *in-vitro* culture was performed. We found that among 559 sRNAs present, 34 sRNAs were differentially expressed at all the three-time points compared to *in vitro* (**Figure 1**). Upregulation or downregulation of these sRNAs at all three time points suggested these sRNAs’ possible role to respond to host circumstances.

**Figure 1.**
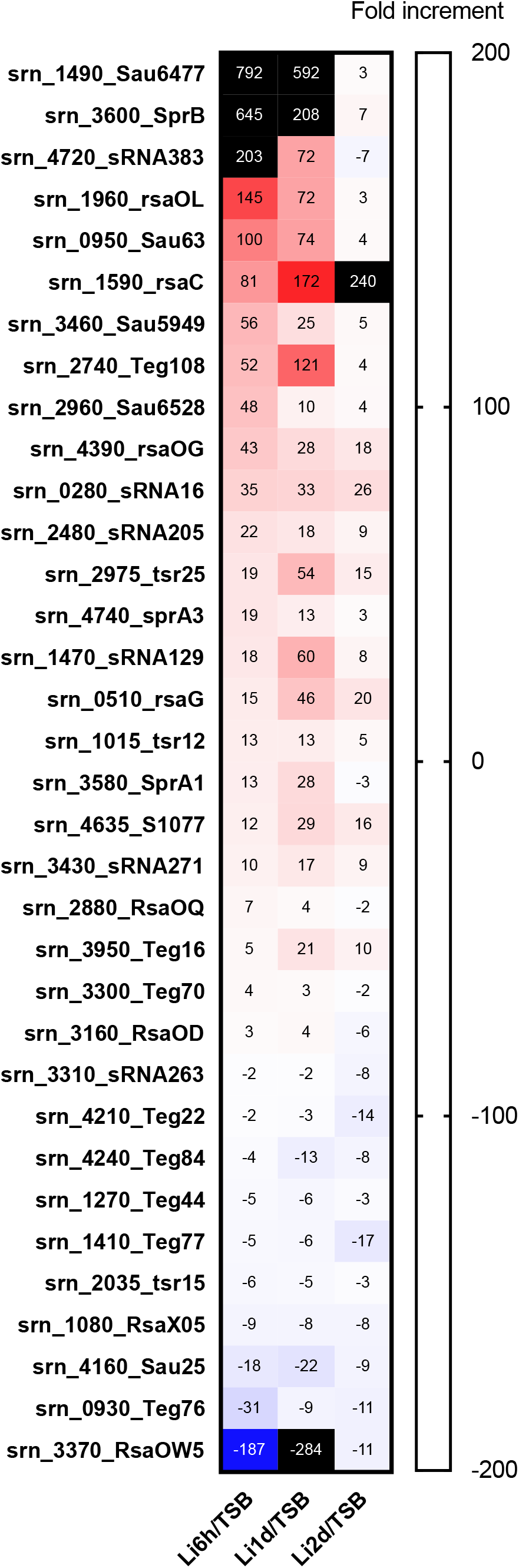
Expression of staphylococcal sRNAs in the host environment. Differential expression of Newman sRNAs in mouse liver compared to TSB medium culture condition. sRNAs with common differential expression and FDR p < 0.05 on all the three-time points are shown.

### Identification of RsaC as a virulence factor

Given that many sRNAs were differentially expressed under the host circumstances, we were interested in whether these sRNAs can modulate *S. aureus* pathogenesis. Among sRNAs, we focused on *rsaC*, as it falls among one of the studied staphylococcal sRNAs. Previously, *rsaC* was shown to have an increased expression in the infected host (15) and modulate oxidative stress during manganese starvation (16). However, its role in the regulation of pathogenesis has remained elusive. First, we confirmed the expression of *rsaC* in mouse organs using a real-time reverse transcription-polymerase chain reaction (RT-PCR). A higher expression of *rsaC* was observed in both the kidney and liver, as compared to TSB medium culture condition (**Figure 2A**). The gene organization of *rsaC* in the Newman genome (**Figure 2B**) showed that *rsaC* did not overlap with other genes, allowing us to construct a single gene disruption mutant. We constructed the *rsaC* disrupted mutant and examined its role in pathogenicity using a mouse infection model. We found that Δ*rsaC* had reduced ability to colonize in mouse heart 24h post-infection whereas the ability to colonize in kidney, liver, lungs and spleen were indistinguishable from that of the wild-type (**Figure 2C**). To further confirm the role of *rsaC* in virulence, we checked the survival of wild-type or Δ*rsaC*- infected mice. The results indicated that the Δ*rsaC* strain had reduced virulence (**Figure 2D**), evident by prolonged survival of the Δ*rsaC* infected mice. To eliminate the possibilities of polar effects associated with this disruption on pathogenicity, we complemented the wild-type and the Δ*rsaC* strains by reintroducing *rsaC* under the control of a constitutive promoter- P*fbaA*. Restored pathogenicity of the complemented strain unequivocally explained the involvement of *rsaC* in pathogenicity and established it as a virulence factor of *S. aureus*.

**Figure 2.**
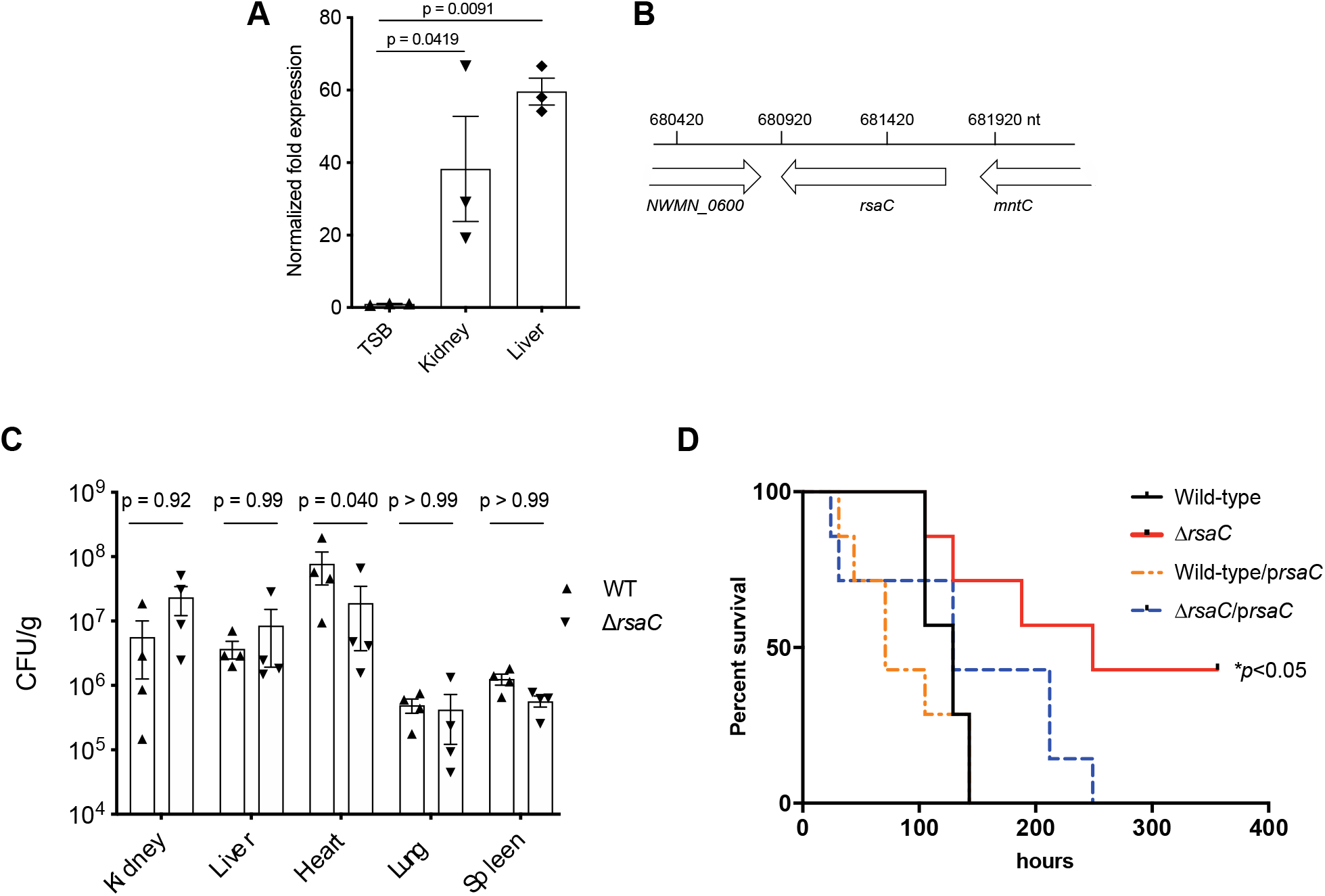
*In vivo* expression and involvement of RsaC in the pathogenesis of *S. aureus*. (A) Upregulation of *rsaC* in host organs compared with that in TSB medium was confirmed by RT-PCR. The data were standardized by the abundance of 16s rRNA in each sample, and statistical analysis was performed using ANOVA. (B) Position of the *rsaC* in *S. aureus* Newman chromosome. (C) The number of *S. aureus* cells in each organ after infection with (7.6×10^7^ CFU and 7.5×10^7^ CFU) *S. aureus* Newman and Δ*rsaC* strains, respectively. Statistical analysis was performed using 2way ANOVA with Sidak’s multiple comparisons in GraphPad Prism 8.4.3. (D) Survival of mice after the infection with a deletion mutant of the *rsaC* gene. Wild-type, Δ*rsaC*, wild-type/p*rsaC*, or Δ*rsaC*/p*rsaC* were injected intravenously at a dose of 5.0×10^7^, 5.9×10^7^, 4.8×10^7^ and 5.0×10^7^ CFU, respectively. Asterisk indicates a significant difference compared with the survival curve following wild-type injection (*p*<0.05) by the Log-rank (Mantel-Cox) test. The experiment was performed two times, and essentially the same results were obtained.

Next, we performed the phenotypic evaluation of the Δ*rsaC* mutant *in vitro*. We found that the growth of Δ*rsaC* strain was indistinguishable from that of the wild-type during both aerobic (**Figure 3A)** and anaerobic **(Figure 3B**) growth. In addition, Δ*rsaC* had unaltered proteolytic and hemolytic abilities compared to that of the wild-type during both aerobic and anaerobic culture conditions (**Figure 3C, D**). These results suggested no difference between the wild-type and mutant in terms of virulence-related *in vitro* phenotypes and necessitated an *in vivo* infection system to investigate the role of *rsaC* in pathogenesis.

**Figure 3.**
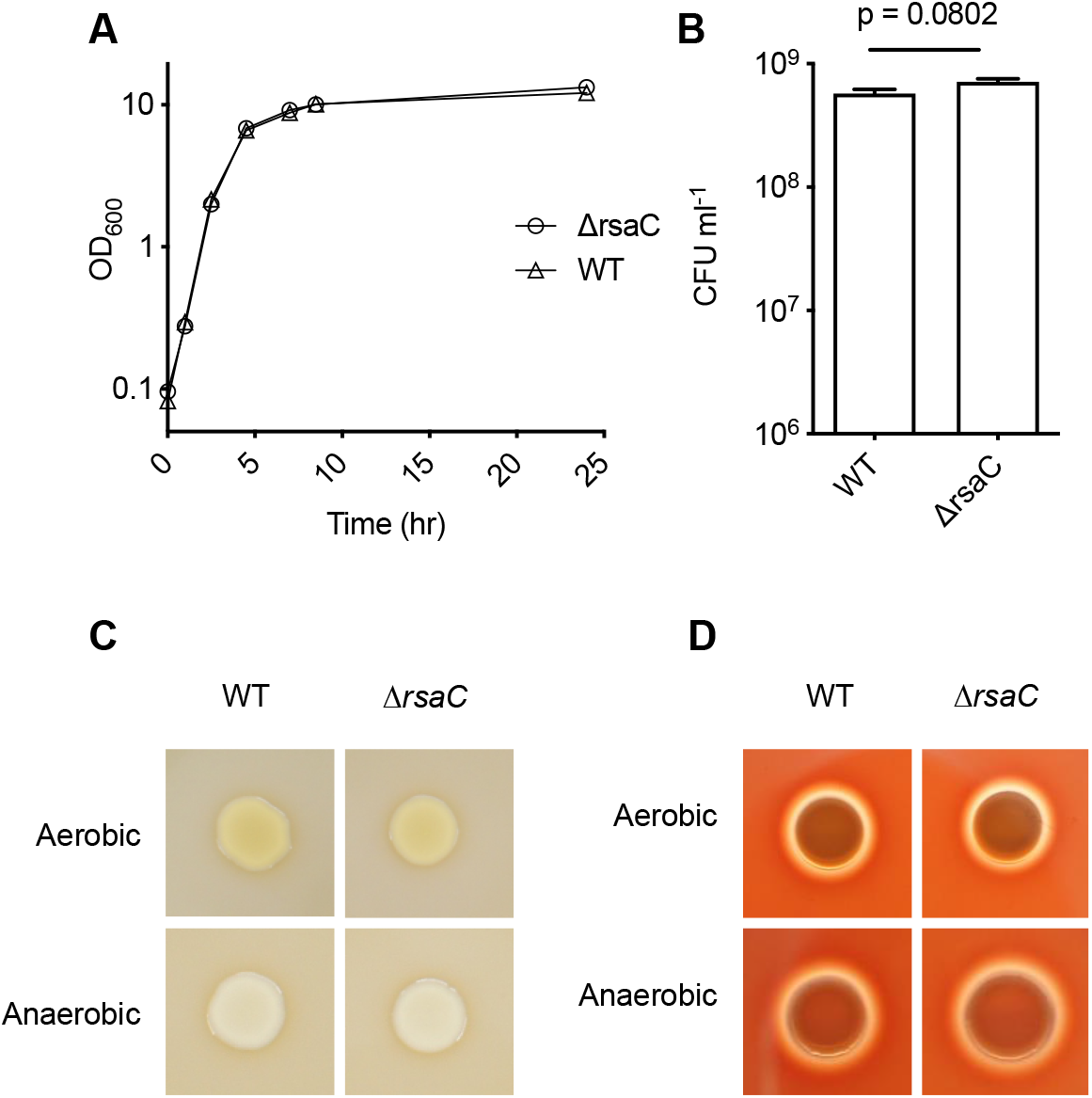
*In vitro* phenotypic analysis of *rsaC* disruption. (A) Growth curve of Δ*rsaC* and wild-type strains during aerobic culture. (B) Colony Forming Units (CFUs) of Δ*rsaC* and wild-type strains after growing anaerobically. Statistical analysis was performed using an unpaired t-test. (C) Proteolytic- and (D) hemolytic- ability of Δ*rsaC* and wild-type strains during aerobic and anaerobic conditions.

### Role of *rsaC* in *S. aureus* transcriptome

Given that *in vitro* phenotypic analysis could not elucidate the possible link between the *rsaC* disruption and pathogenicity, we aimed to examine the transcriptome at the global level. We compared the gene expression pattern by performing an RNA-Seq analysis of the Δ*rsaC* and the wild-type strains under multiple growth conditions- aerobic, anaerobic, and *in vivo* (**Figure 4A**). Since we found higher colonization of *S. aureus* in the heart compared to other organs, and the Δ*rsaC* strain tended to colonize less in the heart compared to the wild-type (**Figure 2D**), we selected heart for *in vivo* RNA-Seq. The gene expression patterns of the Δ*rsaC* strain was compared to that of the wild-type strain in different growth conditions revealing the differential expression of diverse genes in the three tested conditions. Whereas 29, 254, and 47 genes were differentially expressed on RNA obtained from aerobic-growth, anaerobic-growth, and mouse heart, respectively, no commonly affected genes were observed (**Figure 4B**). This indicated a difference in the gene expression pattern depending on growth conditions. Fermentation-related genes, most of which are also associated with the oxidoreductase activity, were downregulated in the Δ*rsaC* mutant when grown under aerobic conditions (**Figure 4C, Table 2**). Based on this, we speculated that *rsaC* directs *S. aureus* towards fermentative respiration by acting as a regulator of fermentation. Under anaerobic conditions, we found the downregulation of many transporters, virulence-related genes in Δ*rsaC* mutant (**Figure 4D, Table 2**). The *in vivo* RNA-Seq analysis, performed in mouse heart at 24 -hr post-infection, showed that compared to wild-type strains, genes related to metal ion (copper and potassium) acquisition and virulence (related to tissue invasion, evasion of host defense, and inhibition of neutrophil activation) were significantly downregulated in the Δ*rsaC* strain (**Figure 4E, Table 2**), which suggested the possible role of host stress towards pathogen response. Taken together, these results indicate the diverse functions of *rsaC* during aerobic, anaerobic, and *in vivo* growth.

**Figure 4.**
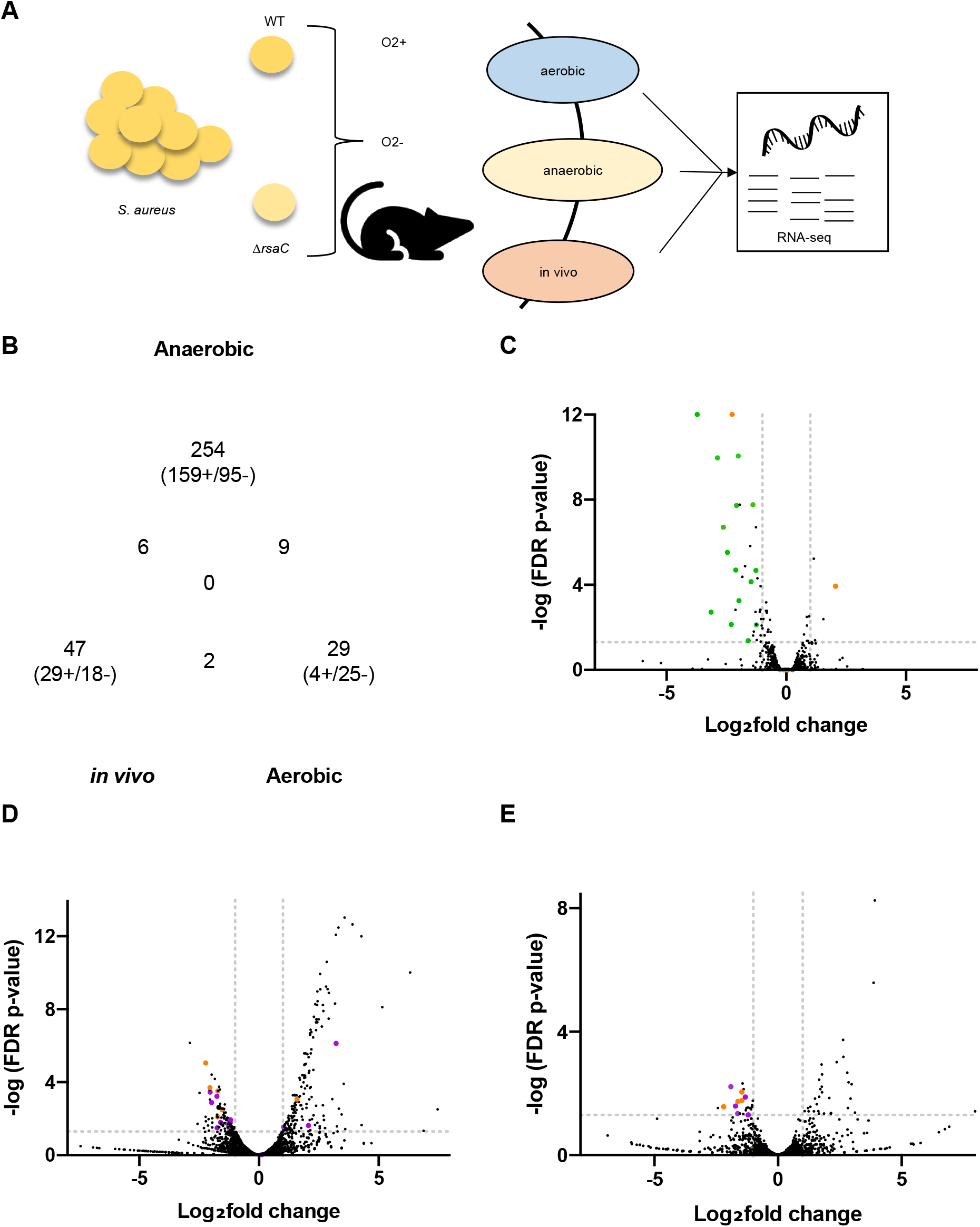
Global transcriptomic analysis of rsaC disruption in *S. aureus*. **(A)** Experimental model to perform transcriptomic analysis. **(B)** Venn diagram showing the genes with common differential expression between *in vitro* (aerobic and anaerobic) and *in vivo* (heart). A volcano plot showing the genes differentially expressed in the Δ*rsaC* strain, compared to wild-type during **(C)** aerobic, **(D)** anaerobic, and **(E)** *in vivo* (heart) growth. Genes involved in fermentation, virulence, and metal-ion acquisition are colored green, orange, and purple, respectively. Data points outside the dotted lines along the y-axis represent fold changes of ≥ 2.0 and that along the x-axis represent FDR *p* ≤ 0.05. The complete list of differentially expressed genes is presented in the supplementary dataset.

**Table 2:**
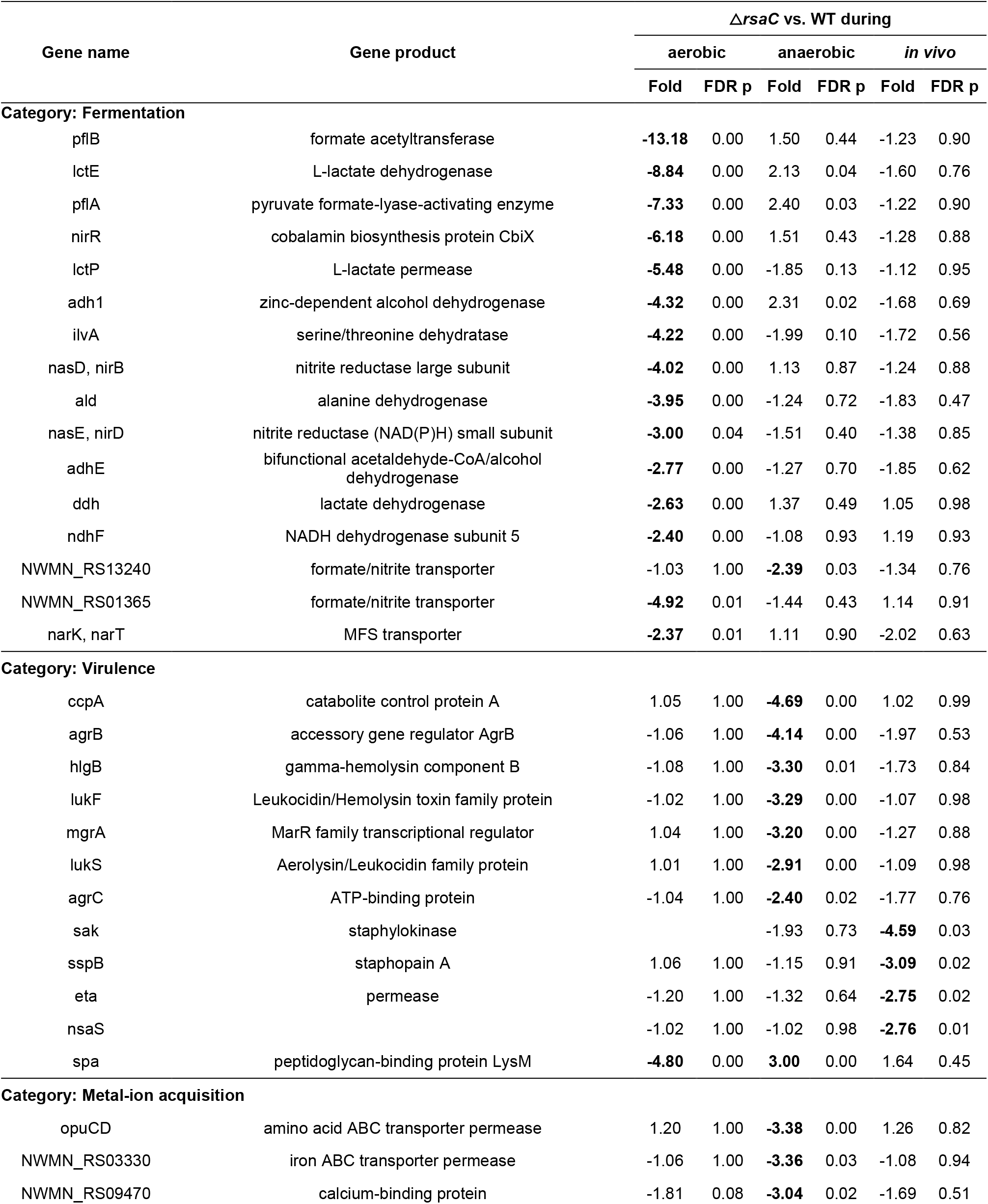
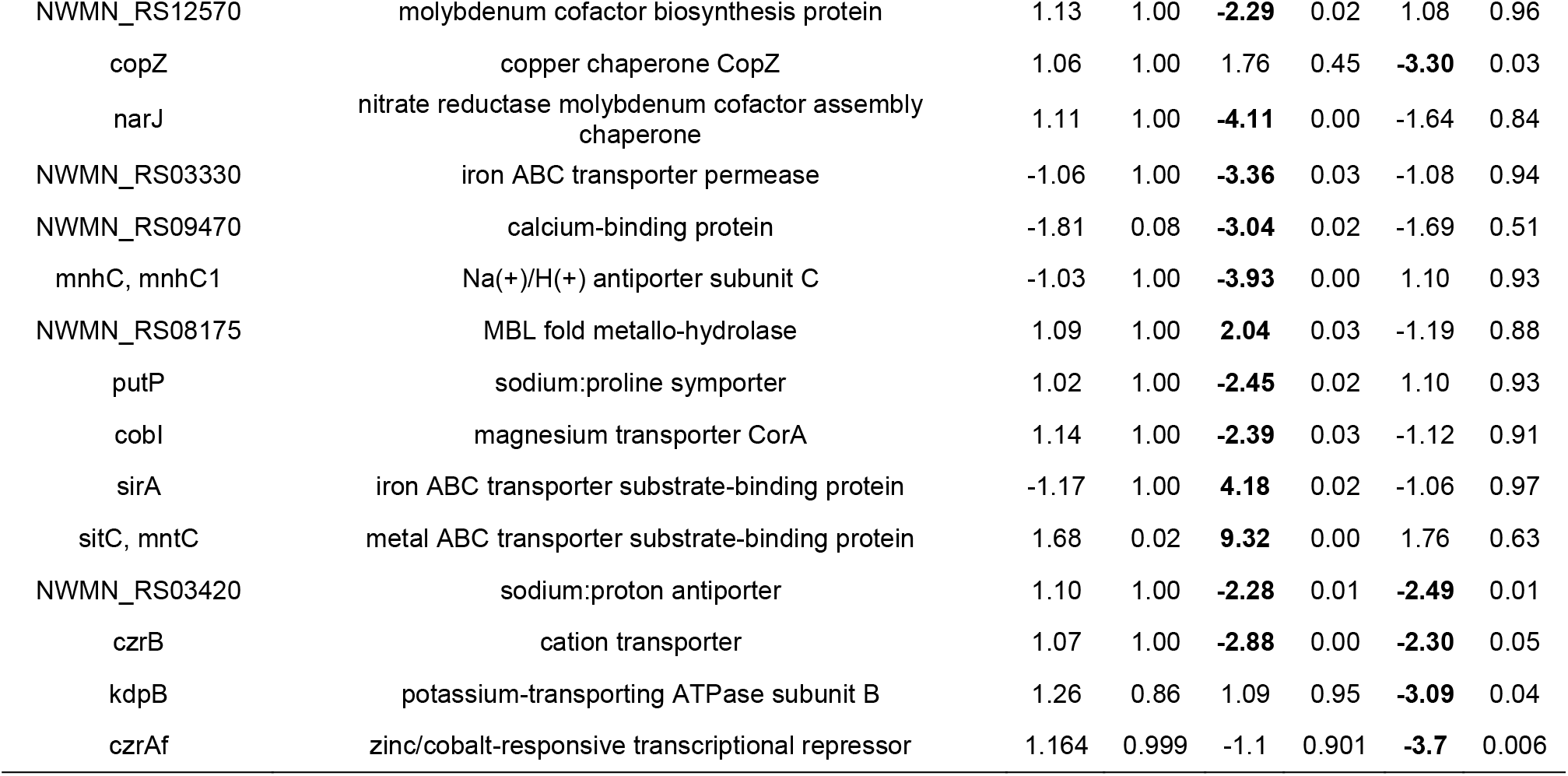
Fermentation, virulence, and metal ion acquisition-related genes differentially expressed (FDR p < 0.05 in the Δ*rsaC* strain compared to wild-type) during aerobic, anaerobic, and *in vivo* growth.

We further categorized the differentially expressed genes depending on their up-and down-regulated status and performed a PANTHER overrepresentation test. We found that genes related to oxidoreductase activity were overrepresented among the genes downregulated in aerobic conditions (**Table 3**). Conversely, among the genes upregulated during the anaerobic condition, the genes related to protein folding and translation were overrepresented, and the genes involved in transmembrane transport were underrepresented (**Table 3**). A previous study also found that these genes are upregulated in anaerobic conditions (26).

**Table 3:**
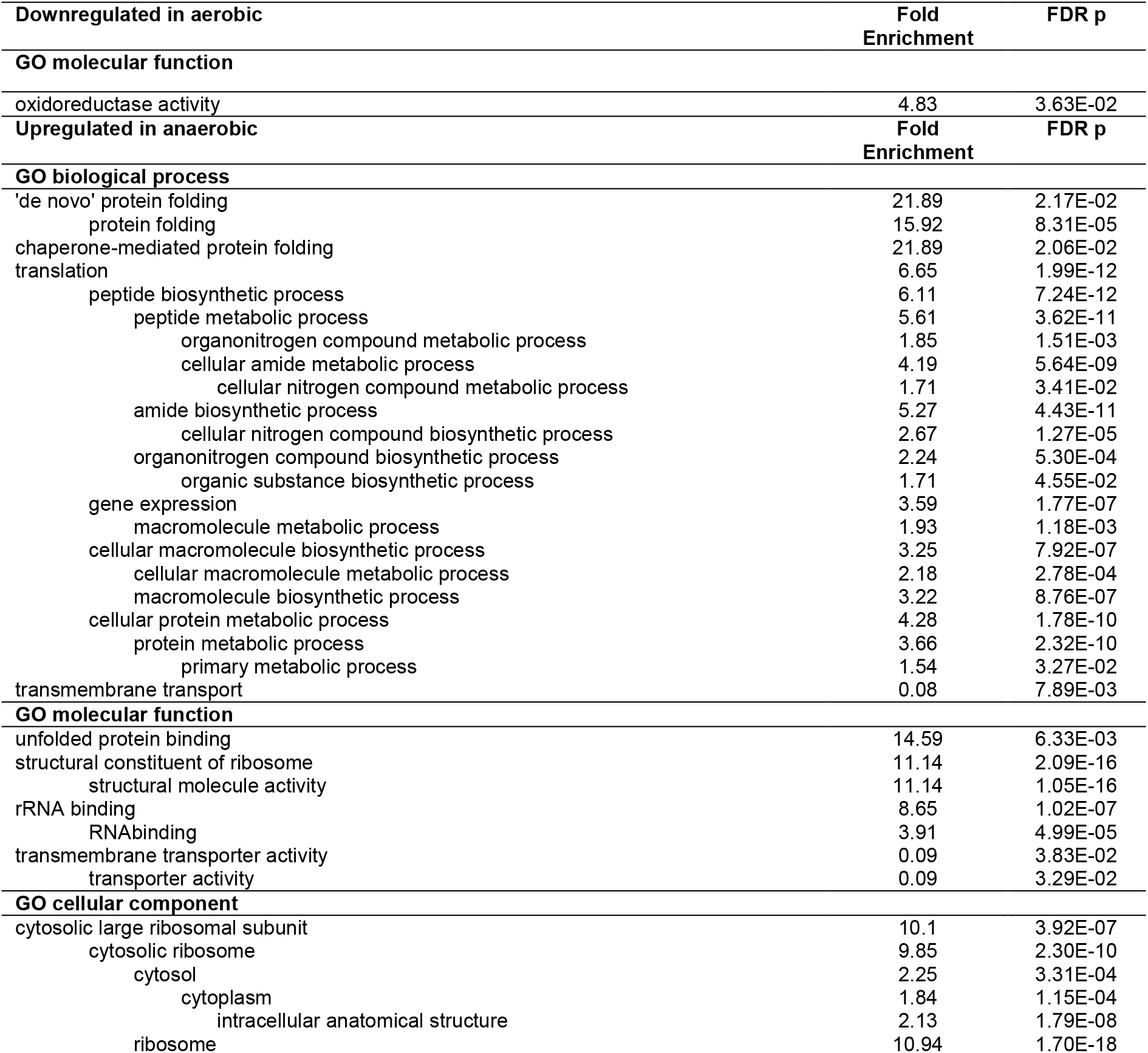
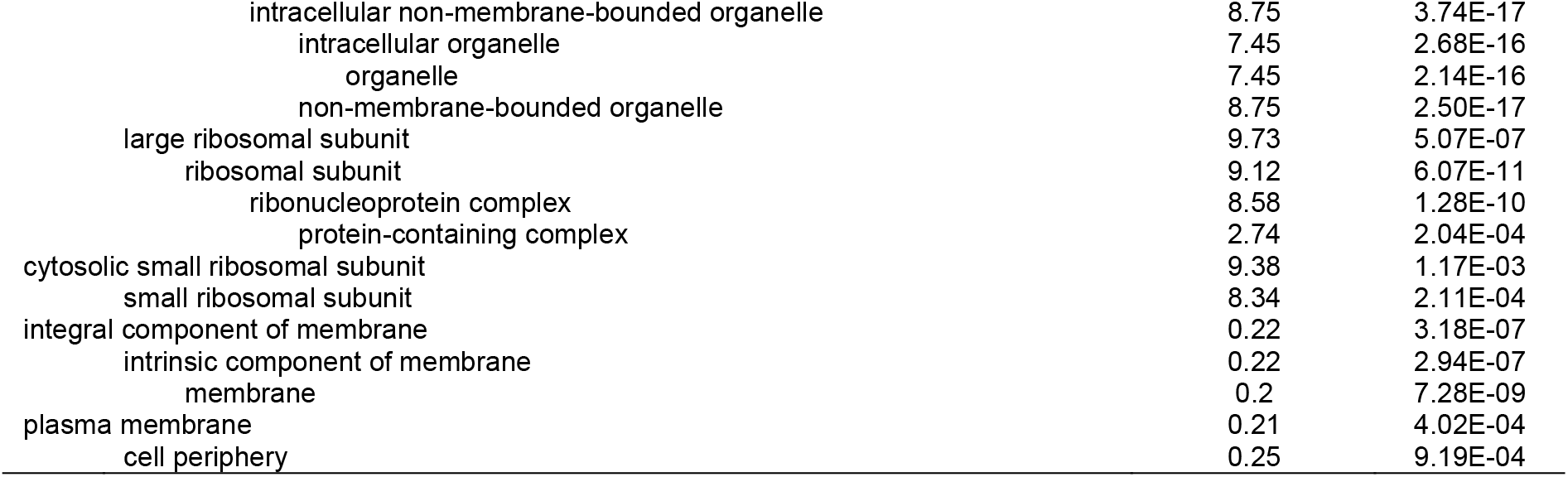
Significantly overrepresented or underrepresented groups among the genes differentially expressed in the Δ*rsaC* strain.

## Discussion

In this manuscript, we identified a Staphylococcal sRNA *rsaC* as a virulence factor, required for full virulence of *S. aureus*. Despite some studies involving *rsaC* (15, 16), its participation in pathogenesis was obscure. Our results suggested that RsaC regulates *S. aureus* transcriptome differently, depending on oxygen availability and host stress. For instance, during aerobic conditions where cells tend to use O_2_ as a terminal electron acceptor (27, 28), we found that RsaC was directing the cells towards anaerobic respiration. Besides, anaerobic conditions, such as those encountered *in vivo* and exaggerated by pathogen colonization and heart failure (6) might lead to the adaptation to adverse environment by regulating nutrition acquisition and virulence factor production, at least partly by the sRNA *rsaC*.

We also found that many genes involved in metal acquisition were downregulated during anaerobic and *in vivo* conditions. Lalaouna et al elucidated the role of *rsaC* in regulating oxidative stress during manganese starvation and identified an Mn-dependent superoxide dismutase *sodA* mRNA as the target of *rsaC* (16). They also found that RsaC interacts of *sodA* mRNA and affects the posttranslational events (16). In our RNA-Seq analysis, we did not find the differential expression of *sodA* in any of the three different growth conditions tested, indicating that *rsaC* does not affect *sodA* at transcriptional level. Thus, our approach of comparing less-virulent strain with its counter virulent strain under different conditions led to the findings that *rsaC* has diverse roles dependent upon the environment encountered by *S. aureus*, yet there existed some commonalities. The gene expression in aerobic condition implied that *rsaC* was involved in shifting bacterial cells from aerobic to anaerobic respiration state; that in anaerobic condition revealed that *rsaC* was involved in metal acquisition and toxin production; and that during *in vivo* condition uncovered that *rsaC* was involved in virulence through host invasion and inhibiting the activation of neutrophils. Therefore, based on previous study (16) and this study, RsaC functions as a transcriptional and translational regulator of a wide variety of genes (**Figure 5**) which ultimately play a role in virulence, a detailed mechanism of which is yet to be identified.

**Figure 5.**
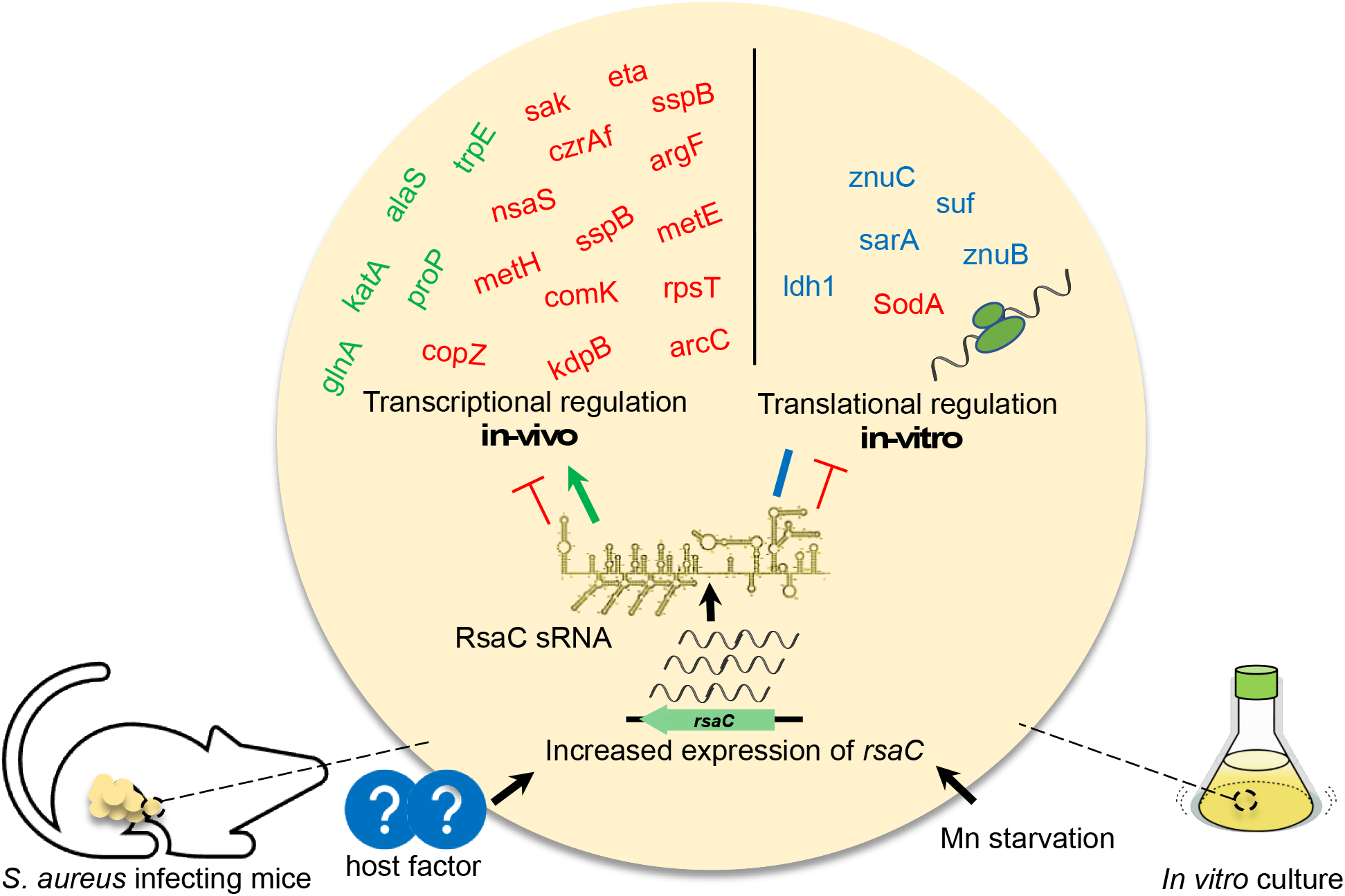
Role of RsaC in gene regulation *in vivo* and *in vitro*. Unknown host factors (*in vivo*) or Mn starvation (*in vitro*) trigger the expression of *rsaC*, which regulates the expression of several genes (this study) or binds to several mRNAs and regulate their translation (16). Red, green, and blue color indicate negative regulation, positive regulation, and binding with unknown effect, respectively.

Further, we observed the upregulation of genes involved in translation, protein folding, and oxidative stress during anaerobic condition. Since clear role of these genes in virulence remains unknown, we speculate that the regulation of translation and protein folding could be a secondary effect of *rsaC* disruption. For instance, in manganese depleted environment, *rsaC* modulated SodM that can use iron as the cofactor for ROS detoxification (16). A detailed understanding of *rsaC* upregulation inside hosts and how RsaC controls gene expression at transcriptional, translational, and posttranslational levels requires further investigation. We searched for partially homologous sequences to the *rsaC* gene in whole genome sequence of the Newman strain that could be targets of transcriptional regulation by RsaC, however; were unable to find any regions. Nonetheless, this key finding of the involvement of RsaC in the virulence of *S. aureus* led to the identification of novel virulence sRNA and opened up avenues for the development of novel antimicrobial agents to treat severe systemic infection by targeting the RsaC signaling pathway.

## Acknowledgements

This work was supported by JSPS KAKENHI Grant Numbers 19K07140JP, 15H05783, 26102714, 24689008, the Mochida Memorial Foundation for Medical and Pharmaceutical Research, and the Takeda Science Foundation to H.H., and in part by JSPS KAKENHI Grant Numbers 19K16653, 20K16253, 21H02733, and JP17F17421, TBRF, IFO fellowships to S.P. and K.S.

## Author contributions

S.P. and H.H. conceived the idea. S.P., H.H. and A.P. performed *in vivo* RNA-Seq analysis. S.P. and A.P. wrote the manuscript. S.O. prepared the gene disruptant mutants. H.H., S.P., A.P., and S.O. performed the mouse systemic infection assays. S.P. performed real time RT-PCR and *in vitro* phenotypic analysis. K.S. critically revised the article for important intellectual content and provided final approval of the article.

